# Hibernation-Driven Immune Suppression in Thirteen-Lined Ground Squirrels

**DOI:** 10.64898/2026.09.14.750722

**Authors:** Miaoyun Zhao, Jackson Chen, Rachel Burrett, Weilong Yang, Subhra Mandal, Saroj Chandra Lohani, Chi Zhang, Jayamanna Wickramasinghe, Matthew T. Andrews, Qingsheng Li

## Abstract

Torpor in mammals imposes extreme energetic constraints, yet how it reshapes the immune system remains poorly understood. Here we characterized splenic immune remodeling during torpor relative to the pre-hibernation active state in a natural hibernator, the 13-lined ground squirrel, using single-cell RNA sequencing, immunohistochemical staining, and quantitative image analysis. The torpid spleen showed a significantly reduced white-pulp area fraction with a preferential reduction of B cell- and T cell-rich adaptive niches and relative preservation of innate myeloid populations. Among splenic immune cells, the relative frequency of B cells was significantly lower and that of macrophages significantly higher in torpid animals. Across five immune cell types examined (B cells, T cells, macrophages, dendritic cells, and neutrophils), torpid animals exhibited more than 2-fold higher *Cirbp* expression (nominal *P* < 0.05); *Txnip* was also higher and *Hk2* lower, with adjusted *P* < 0.05 in B and T cells, suggesting reduced glycolysis in these cells. In B and T cells, canonical pathway analysis predicted inhibition of mitogen-activated protein kinase, mTOR, and chemokine signaling, together with activation of capped intron-containing pre-mRNA processing. In myeloid cells, predicted pathway changes differed by cell type, including inhibition of the RHO GTPase cycle in macrophages and of inflammasome signaling in dendritic cells, and activation of interleukin-1 family signaling in neutrophils. These findings identify selective splenic immune remodeling associated with torpor, characterized by a reduced white-pulp area fraction and altered immune cell composition, accompanied by shared changes in *Cirbp*, *Txnip*, and *Hk2* expression and cell-type-specific transcriptional responses.

## Introduction

The immune system is vital for maintaining organismal homeostasis and survival under stressful or threatening conditions^1,2^. Immune dysregulation contributes to the pathogenesis of infectious diseases, cancer, and autoimmune diseases^3,4^. In chronic infection and cancer, persistent antigen and inflammatory signals promote T cell exhaustion and impaired effector function^3^. Conversely, autoimmune diseases result from failed immune regulation and loss of self-tolerance^5^.

Hibernation is a seasonal physiological state characterized by prolonged bouts of torpor, during which metabolic rate and body temperature are markedly reduced^6^. Studies of mammalian hibernation have inspired a growing field of hibernation-based medicine, including enhancement of organ preservation for transplantation, mitigation of traumatic brain injuries and heart attack, management of hemorrhagic shock, and astronaut radiation protection in space exploration^6–10^. However, the immune regulatory mechanisms that govern the transition between torpor and active state remain poorly understood.

Hibernation in 13-lined ground squirrels (*Ictidomys tridecemlineatus*; hereafter 13-LGS) represents a naturally evolved state of profound metabolic depression that supports prolonged fasting and adapts to low environmental temperature^11–14^. During hibernation, 13-LGS remain underground for about 6–7 months, from early autumn to spring, alternating between extended torpor bouts (7–15 days) and short interbout arousals (IBA, 12–18 h)^6,15,16^. During torpor, their body temperature (*T*_b_) falls to near ambient temperature, heart rate to 3–5 beats per minute, and respiratory rate decreases to only a few breaths per minute^6,17,18^. In summer, 13-LGS are in an active, non-hibernating state characterized by normal feeding, locomotor activity, and physiological homeostasis. Physiological adaptations that occur during torpor inherently constrain the immune system, as effective host defense depends on metabolically demanding processes such as cytokine production, immune cell migration, and proliferation^19^. Multiple studies have demonstrated that torpor is associated with reduced responsiveness in both innate and adaptive immunity^20,21^. A defining feature of immune suppression during torpor is a profound yet reversible peripheral blood leukopenia, in which circulating white blood cell counts can decline by more than 90%^22,23^. In addition, classical pathway serum complement activity is significantly reduced during torpor although it remains functional *in vitro* even at 5 °C^24^. Experimental evidence further supports immune suppression during torpor: torpid golden-mantled ground squirrels failed to exhibit a typical febrile response following lipopolysaccharide (LPS) injection^20^, hibernating 13-LGS did not generate antibody responses to T cell–independent antigens^25^, and the onset of skin allograft rejection was considerably delayed^26^, indicating diminished humoral and cellular immune responses. Together, these findings underscore a state of coordinated immune suppression during torpor.

However, most hibernation-related immunological studies have primarily examined immune cells from peripheral blood, which represents only about 2% of the body’s total immune cells^27^. Consequently, analyses restricted to blood cells cannot fully capture how immune niches are reorganized or how lineage-specific immune cells are regulated during torpor. Secondary lymphoid tissues (LTs), such as the spleen, perform critical immune functions, where immune cells reside and are maintained. Although it is the largest lymphoid organ, the spleen remains understudied in relation to immune adaptation during hibernation. Structurally, the spleen integrates anatomically distinct yet functionally coupled compartments. The red pulp removes senescent erythrocytes and serves as a storage site for platelets, leukocytes and red blood cells, while the white-pulp is enriched in immune cell populations and is the major architecture underpinning adaptive immunity^28,29^. This unique organization positions the spleen as a critical hub for orchestrating immune cell migration and functional modulation throughout torpor. However, the molecular mechanisms governing these adaptations within LTs remain poorly understood, and the immune cell-type–specific transcriptomic responses to torpor compared to the active state have yet to be characterized. To address this gap, we reasoned that the comparison of torpid with pre-hibernation active animals provides a useful framework for defining the immune adaptations associated with torpor. We combined quantitative image analysis (QIA) of immune cell populations in splenic tissues with single-cell RNA sequencing (scRNA-seq) of splenic immune cells from active and torpid 13-LGS to delineate structural, compositional, and lineage-specific transcriptional alterations underlying immune adaptation during torpor.

## Materials and methods

### Study design and ethics statement

The animal study was approved by the Institutional Animal Care and Use Committee (IACUC) at the University of Nebraska-Lincoln (UNL) (Protocol # 2374) and conducted as reported previously^30^. Briefly, 13-LGS were captured near Lincoln, Nebraska (July–August) and singly housed at ∼20 °C with ad libitum food and water under a 12:12 h light/dark cycle at UNL veterinary facilities under the care of UNL Institutional Animal Care Program veterinary staff. On October 1, room temperature was decreased to 12 °C to facilitate the entry into torpor. Beginning in early November, squirrels were transferred to an environmental chamber (5 °C, constant darkness, no food; water available), conditions that reliably induce deep torpor. To assess hibernation-associated changes in the spleen, animals were euthanized at torpid and active states: (i) early September, an active pre-hibernation state (*T*_b_ ∼35 °C) (*n* = 7), and (ii) December–January, during deep torpor (*T*_b_ 6.2–7.2 °C) (*n* = 8). Following euthanasia, the spleen was dissected out, and its weight and dimension were measured. A small portion of spleen was fixed in SafeFix II All-Purpose Fixative (Cat. #23-042600, Fisher Scientific, MA) and embedded into a paraffin block for tissue analysis and fresh tissues were used for immune cell isolation and scRNA-seq as described below.

### Immunohistochemical staining and quantitative image analysis of immune cells in spleen

The fixed splenic tissues were systematically divided into three anatomically distinct portions of proximal, central, and distal, providing representative sampling coverage across the full extent of the spleen. Tissue sections of 5 µm from each portion were subjected to Hematoxylin and Eosin (H&E) and immunohistochemical staining (IHCS) using antibodies against various immune-cell markers. For H&E staining, after deparaffinization and rehydration, tissue sections on slides were stained with 50% hematoxylin (Cat. #3530-1; Ricca Chemical, Arlington, TX, USA) for 1 min and rinsed in distilled water. The sections were then counterstained with Edgar Degas Eosin solution (Cat. #HTE-GL; BioCare Medical, Pacheco, CA) for 3 min, followed by dehydration through graded ethanol and clearing in xylene. For IHCS, tissue sections on slides were subjected to antigen retrieval in 1× citrate target retrieval buffer (pH 6.0) using heat-mediated retrieval for 15 min. Endogenous peroxidase activity was quenched with 3% hydrogen peroxide for 10 min at room temperature, followed by blocking with 5% bovine serum albumin in 1× TBST for 1 h at room temperature. Sections were incubated overnight at 4 °C with primary antibodies against CD21 (1:500 dilution, Cat. No. ab227662, Abcam), CD3 (1:200 dilution, Cat. No. ab245731, Abcam), CD4 (1:200 dilution, Cat. No. ab133616, Abcam), and Ki67 (1:200 dilution, Cat. No. 275R-14, Cell Marque). After washing with 1× TBST, the sections were incubated with an anti-rabbit polymer (Cat. No. K4003, Dako) and developed using 3,3′-diaminobenzidine (DAB) substrate. Slides were counterstained with hematoxylin, dehydrated through graded ethanol, cleared in xylene, and mounted. Whole-slide images were scanned using a Leica MICA imaging system. To quantify DAB-immunochemical stained immune cells or densely nucleated white-pulp in H&E–stained sections, digitized whole-slide images were analyzed using QuPath software (version 0.6.0) according to the published methods with modifications^31^. Briefly, the tissue region was manually delineated to define the total analyzable tissue area and exclude non-tissue background, glass-slide artifacts, tissue folds, tears, and other regions unsuitable for analysis. Regions of white-pulp where there are densely nucleated cells in H&E sections were selected consistently across all specimens. To separate the DAB-chromogen signal from the hematoxylin counterstain and background signals, the whole tissue images were subjected to color deconvolution. DAB-positive areas were identified using an intensity threshold. The resulting positive-area annotations were visually reviewed to verify accurate discrimination of specific DAB staining from background signal and nonspecific staining. Results were reported as DAB-positive area divided by total analyzed tissue area (%). The marker-positive area fraction was used as an index of the relative abundance of marker-positive cells.

### Peanut agglutinin fluorescence staining

After deparaffinization and rehydration, splenic tissue sections were incubated with neuraminidase (0.1 U/mL; Cat. No. N2876, Sigma-Aldrich) in 50 mM sodium acetate buffer containing 5 mM CaCl₂ (pH 5.5) for 45 min at 37 °C to expose PNA-binding glycans. Sections were then washed three times for 5 min each and blocked with 1% bovine serum albumin (BSA) in wash buffer for 30 min at room temperature. Sections were incubated with Alexa Fluor 647-conjugated peanut agglutinin (PNA; Cat. #L32460, Invitrogen) at 10 µg/mL in 1× TBS containing 1% BSA and 5 mM CaCl₂ for 45 min at room temperature in the dark. After incubation, sections were washed three times for 5 min each, counterstained with DAPI, and mounted using an antifade mounting medium. Fluorescence images were acquired using identical imaging settings across experimental groups. PNA staining was evaluated for the presence and distribution of PNA-positive germinal-center B cells in white-pulp.

### Isolation of splenocytes for scRNA-seq

Fresh splenic tissue was processed into single-cell suspension using an adapted mechanical dissociation protocol based on published mouse splenocyte preparation methods and protocols developed for single-cell transcriptomics^32^. Briefly, spleen tissues were mechanically dissociated by gently pressing them through a 70 µm nylon cell strainer using Dulbecco’s phosphate-buffered saline (DPBS; Cat. No. 21-031-CM, Corning, Manassas, VA) supplemented with 2% fetal bovine serum (FBS; HyClone, Rockford, IL), hereafter referred to as isolation buffer. Red blood cells were removed by resuspending the splenocyte pellet in ACK (Ammonium–Chloride–Potassium) lysis buffer and incubating the suspension on ice for 10 min. Following RBC lysis, cells were treated with DNase I (20 µg/mL) for 15 min with gentle mixing to reduce aggregation. The resulting suspension was passed through a 70 µm nylon cell strainer once more to obtain a uniform single-cell preparation for downstream analyses.

### Library preparation and sequencing

scRNA-seq libraries were generated using the Chromium Single Cell 3′ v3 platform (10x Genomics). Libraries were sequenced on an Illumina NovaSeq 6000 S2 flow cell with 2 × 50 bp paired-end reads to a depth of ∼300 million reads per library.

### Alignment, cell clustering, and cell-population annotation

BCL files were converted to FASTQ with Cell Ranger suite (v9.0.0, https://support.10xgenomics.com)^33^. Cell Ranger was also used to align (via STAR(v2.7.9a))^34^ raw sequencing data to the 13-LGS (SpeTri2.0) reference genome downloaded from GenBank (GCA_000236235.1) and quantify UMI counts. Then the UMI counts were loaded to R using Seurat package (v5.3.0)^35^ and integrated. Subsequent quality control steps included filtering out cells with fewer than 500 detected genes in total UMI counts, based on visual inspection of the distributions of these metrics. Cells with a mitochondrial read percentage exceeding 5% were also excluded to remove damaged cells or cells in apoptosis. Doublet detection was performed using scDblFinder (v1.22.0)^36^ with p = 0.3, and predicted doublets were removed from the dataset. Next, the UMI counts were normalized using Seurat’s default normalization method and then scaled. Principal Component Analysis (PCA) was performed on the scaled expression matrix (HVGs only) to reduce the dimensionality, retaining the top 50 principal components. The cells were then embedded into a 2-dimensional space using Uniform Manifold Approximation and Projection (UMAP) for visualization. Cell clustering was performed using the default algorithm, which uses PCA-reduced data by iteratively grouping cells into clusters by optimizing a modularity function, aiming to maximize connections within clusters while minimizing connections between them. The clustering resolution was optimized to identify distinct cell populations based on known marker gene expression and visual inspection of the UMAP embedding. Cell clusters were annotated based on the expression of known marker genes obtained from the ImmGenData (v2024-02-26)^37^ (only with orthologous genes) and celldex (v1.18.0) databases using SingleR R package^38^. Clusters were assigned to specific cell types (e.g., T cells, B cells, macrophages) based on the enriched expression of their respective marker genes. Differential gene expression (bulk RNA obtained with Seurat AggregateExpression) analysis was conducted between cell types (or conditions) using DESeq2^39^. UMAP plots were used to visualize the overall cellular landscape and highlight different cell types between selected cell groups. All visualizations were generated using custom scripts in R leveraging packages like ggplot2^40^.

Differential gene expression between torpid and active animals was analyzed within each cell type using DESeq2 with counts aggregated using Seurat AggregateExpression. *P*-values were adjusted using the Benjamini–Hochberg method for multiple testing. Genes meeting both adjusted *P* < 0.05 and |log₂ fold change| > 1 were considered differentially expressed genes (DEGs). Volcano plots show log₂ fold changes against −log₁₀(nominal *P*). For visualization, genes meeting both nominal *P* < 0.05 and |log₂ fold change| > 2 were highlighted in red. Vertical dashed lines indicate log₂ fold changes of −2 and +2, and the horizontal dashed line indicates nominal *P* = 0.05. Heatmaps display selected genes meeting both adjusted *P* < 0.05 and |log₂ fold change| > 1; for dendritic cells (DCs), all five qualifying genes are shown. Expression values were standardized to row-wise Z-scores across animals. Red and blue indicate expression above and below the mean for each gene, respectively.

Pathway analysis was performed using Ingenuity Pathway Analysis (IPA; QIAGEN, build version 9.0, content version 169997502, release date 2026-08-16), with input identifiers specified as mouse gene symbols. For all cell types, input genes were selected using nominal *P* < 0.05 and a greater than 1.5-fold change in either direction. The reference set was the Ingenuity Knowledge Base (Genes Only). Canonical pathways with Benjamini–Hochberg-adjusted *P* < 0.05 were considered enriched; activation z-scores ≥ 2 or ≤ −2 indicated predicted activation or inhibition, respectively. Twenty representative pathways meeting both criteria were selected for display for each cell type. Selection considered relevance to splenic immune cells, the number of input genes, and overlap in pathway members; displayed pathways are not independent of one another.

### Statistical analysis

Group comparisons using unpaired two-sided Welch’s *t*-test were performed in GraphPad Prism (version 9.4.1). Each animal was treated as an independent biological replicate. Data are presented as mean ± SEM, and *\*P* < 0.05 was considered statistically significant. The statistical tests applied and the number of animals analyzed per group were specified in the corresponding materials and methods sections and figure legends.

## Results

### Torpor Selectively Remodeled Spleen Architecture

It has been previously documented that peripheral blood immune cells undergo a marked reduction (leukopenia) during torpor^22,23^. A prevailing hypothesis suggests that this reduction reflects the migration of circulating immune cells into secondary lymphoid organs, especially the spleen. To test this hypothesis and elucidate how the spleen adapts to the extreme physiological challenge during torpor, we compared splenic size and weight between torpid and active animals at the time of euthanasia. Torpid animals (*n*=8) weighed less than active animals (*n*=7) (165.75 ± 5.11 vs. 216.71 ± 16.12 g; *P* = 0.019). Mean spleen weight was also lower in torpid animals, but the difference was not statistically significant (0.211 ± 0.022 vs. 0.270 ± 0.040 g; *P* = 0.226). Despite the profound metabolic suppression associated with torpor, the splenic index (spleen weight/body weight, *P* = 0.988) as well as spleen length (*P* = 0.853) did not differ significantly between torpid and active 13-LGS (Fig. 1A), which does not support the global splenic atrophy during torpor.

**Figure 1.**
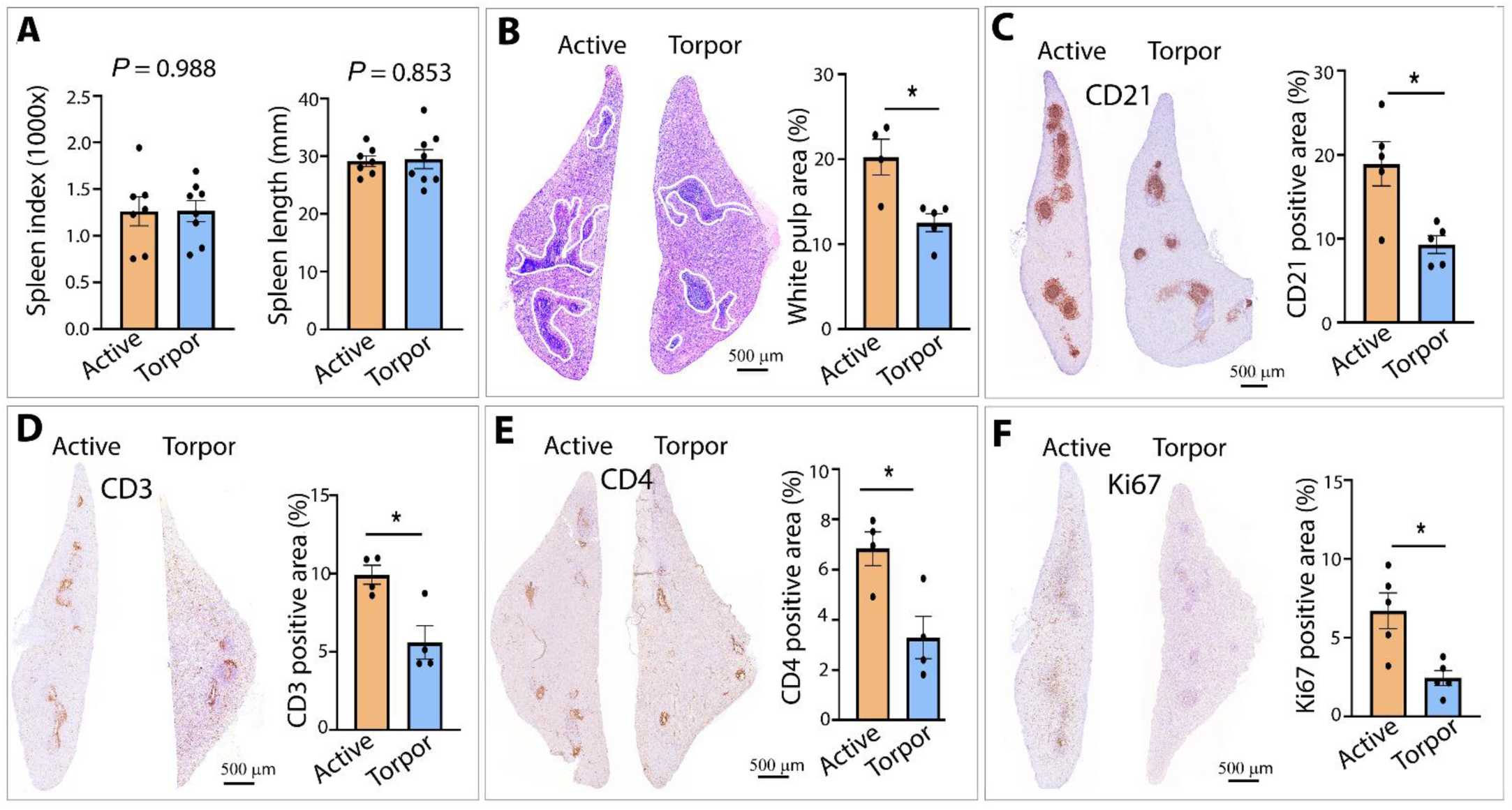
Torpor selectively remodels splenic architecture without significant differences in spleen index or length. (A) Spleen index, defined as spleen weight normalized to body weight and multiplied by 1,000, and spleen length in active (*n* = 7) and torpid (*n* = 8) 13-lined ground squirrels (13-LGS). Exact *P* values are shown. (B) Representative hematoxylin and eosin (H&E)-stained spleen sections from active and torpid animals. White outlines delineate white pulp. Quantitative image analysis (QIA) shows the percentage of total splenic tissue area occupied by white pulp. (**C–F**) Representative immunohistochemical staining (IHCS) and QIA of areas positive for CD21+ cells (**C**), CD3+ T cells (**D**), CD4+ T cells (**E**), and Ki67 (**F**). *n* = 4–5 per group, each point represents one animal. Bars represent mean ± SEM. Statistical comparisons were performed using unpaired two-tailed Welch’s *t*-test. \**P* < 0.05. Scale bars, 500 µm.

Quantitative image analysis (QIA) of H&E-stained splenic sections of torpid and active animals (*n* = 4–5 per group), however, revealed notable yet localized remodeling of splenic architecture between these two groups (Fig. 1B). In particular, the proportion of white-pulp, which is enriched in T cell–dense periarteriolar lymphoid sheaths (PALS) and B cell–rich follicles and collectively serves as secondary lymphatic tissue compartments, was significantly reduced in torpid animals (12.53 ± 1.05%) compared to active controls (20.26 ± 2.10%) (Fig. 1B; *P* < 0.05). We compared germinal-center B cells within the white-pulp by peanut agglutinin (PNA) staining and observed a reduction in PNA+ signals within the white pulp (Supplementary Fig. 1). We further quantified CD21+ cells (several B-cell subsets, including marginal-zone B cells as well as follicular dendritic cells) using a combination of IHCS and QIA (Fig. 1C). The frequency of CD21+ cells was significantly reduced in torpid animals compared with active animals (Fig. 1C). T-cell-associated compartments were also reduced in torpid animals compared with active animals (Fig. 1D&E). The CD3-positive staining-area fraction was lower in torpid animals compared with active animals (Fig. 1D). Similarly, the CD4+ cells in torpid animals were also significantly reduced compared with active animals (Fig. 1E). Together, these findings indicate that torpor-associated remodeling encompasses both B cell- and T cell-associated lymphoid compartments. Consistent with the reduction of abundance of B cells and T cells, Ki67, a proliferation marker, was also significantly reduced during torpor, especially in white-pulp cells (Fig. 1F). This selective contraction of B and T cells and the size of white-pulp indicate a downscaling of the adaptive immune component of the spleen while relatively maintaining red-pulp associated roles in blood filtration and storage^28,41–43^.

Collectively, these results demonstrated that torpor selectively remodeled the spleen without a significant change in spleen mass. This remodeling is characterized by contraction of the white-pulp, reductions in PNA+ B cells, CD21+ cells, CD3+ and CD4+ T cells, and decreased cell proliferation. These coordinated changes are consistent with the reorganization and suppression of splenic adaptive immune arm during torpor.

### scRNA-seq Showed the Different Splenic Immune Cell Landscape in Active and Torpid Groups

Given the pronounced remodeling of immune cell niche, white-pulp during torpor, we next examined how torpor reshapes splenic immune cell composition and transcriptional states by performing scRNA-seq of immune cells isolated from the spleens of active (*n* = 4) and torpid (*n* = 4) 13-LGS (Fig. 2A). Single cells were sequenced using the 10x Genomics platform, and low-quality transcriptomes (<500 genes detected or >5% mitochondrial reads) were removed before downstream analysis. After passing quality control, a total of 46,685 immune cell transcriptomes were retained, comprising 21,341 immune cells from active animals and 25,344 immune cells from torpid animals. Integrated analysis using Seurat revealed 16 transcriptionally distinct clusters visualized by UMAP (Fig. 2B). Cluster annotation using both mouse ortholog mapping and ImmGen reference datasets identified all major splenic immune cell populations, including but not limited to B cells, T cells, macrophages, DCs, natural killer (NK) cells and natural killer T (NKT) cells, eosinophils, and neutrophils (Fig. 2C). Splenic immune cells from active and torpid animals clearly differed in their distribution across the shared UMAP embedding, with several regions more densely populated in torpid animals (Fig. 2C).

**Figure 2.**
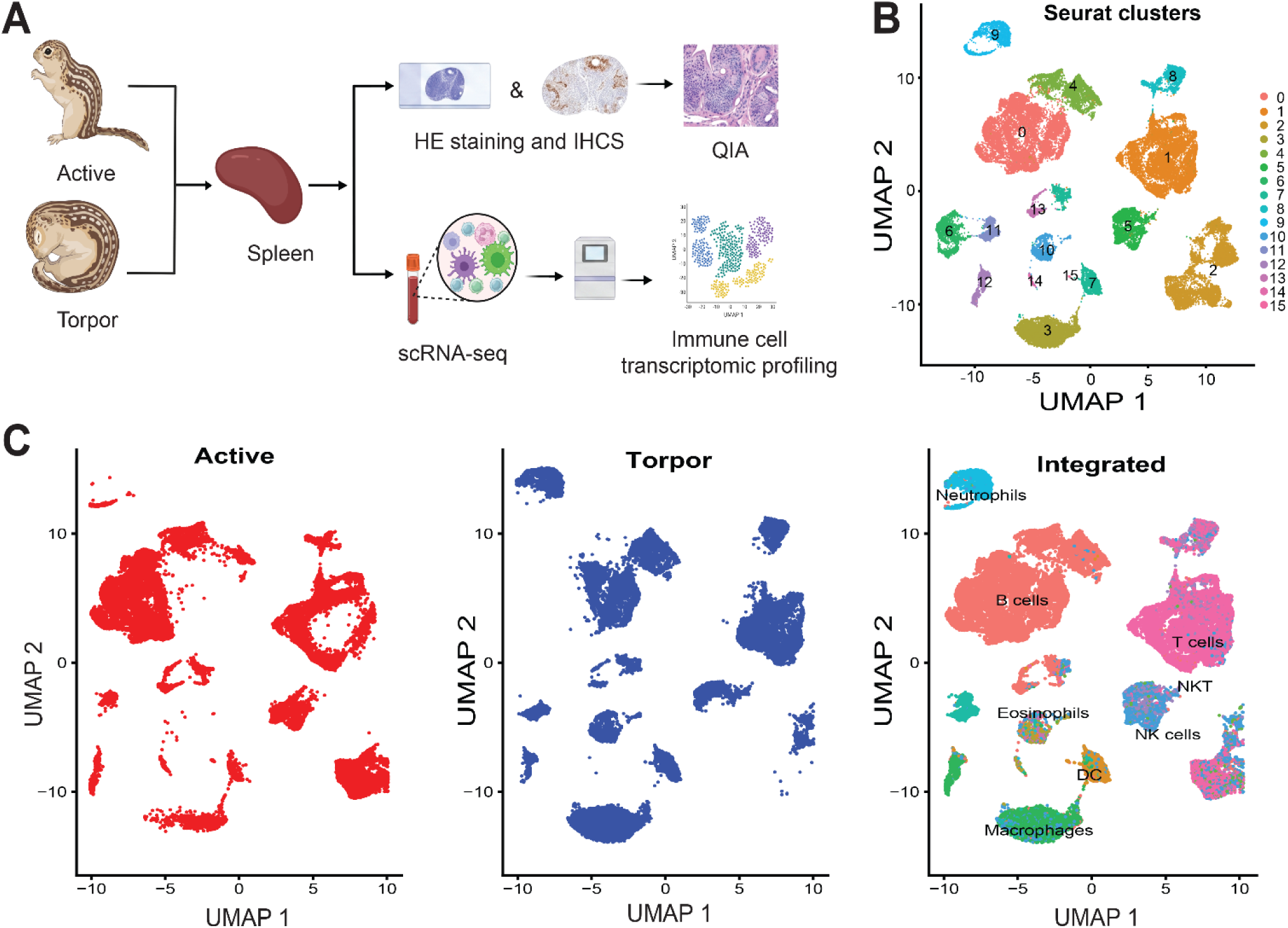
Single-cell RNA sequencing delineates splenic immune cell composition in active and torpid animals. (**A**) Schematic overview of the histological, immunohistochemical, quantitative image analysis, and single-cell RNA sequencing workflows. Tissue sections were subjected to hematoxylin and eosin (H&E) staining, immunohistochemical staining (IHCS), and quantitative image analysis (QIA). Spleens were collected from active (n = 4) and torpor (n = 4) animals, dissociated into single-cell suspensions, processed using the 10x Genomics Chromium platform, and subjected to downstream bioinformatic analysis. Panel A was created with BioRender.com. (**B**) Uniform manifold approximation and projection (UMAP) visualization of the integrated splenic single-cell dataset following Seurat-based clustering. Transcriptionally distinct cell clusters are shown and numbered. (**C**) UMAP representation of the integrated dataset annotated by major immune cell populations (right), with cells originating from active animals (left, red) and torpor animals (middle, blue) projected onto the same UMAP embedding to visualize their relative distribution across immune cell populations.

To assess potential dissociation-induced artifacts, we examined expression of canonical immediate-early and stress-response genes (*Fos, Jun, Dusp1, Egr1, Atf3*, and *Hspa8*) across immune lineages between torpid and active animals. These markers did not exhibit a global induction pattern, and *Fos* remained unchanged in the four cell types with available adjusted *P* values (Supplementary Table 1), indicating that torpor-associated transcriptional differences are unlikely to be driven by handling-induced stress.

### Reduced Abundance and Altered Transcriptional Profile of Splenic B Cells during Torpor

Consistent with the observed reduction of B cells in white-pulp during torpor using CD21 IHCS (Fig. 1C) as well as PNA staining (Supplementary Fig. 1), one of the most striking observations from our scRNA-seq data was the significant decrease in the relative abundance of B cells during torpor, with B cells comprising 44.3 ± 3.6% of total immune cells in active animals but declining to 21.1 ± 3.3% in torpid animals, with a 52.4% reduction (*P* < 0.05) (Fig. 3A). In addition to the reduced B-cell abundance, UMAP visualization showed that B cells from torpid animals formed a discrete and transcriptionally distinct cluster from that of active animals, indicating a global alteration in B cell transcriptional program rather than solely population reduction (Fig. 3B).

**Figure 3.**
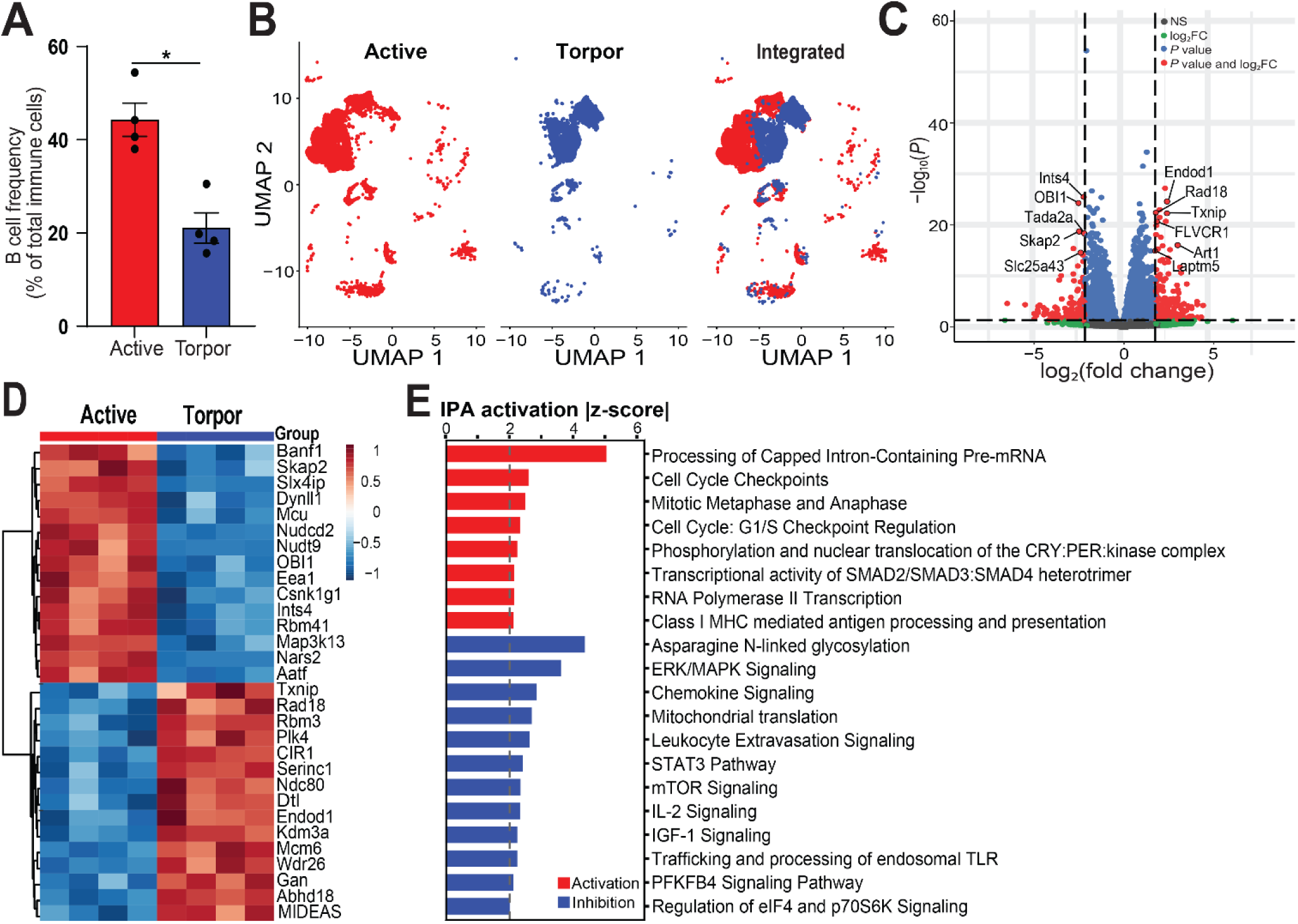
Torpor alters the abundance and transcriptional profile of splenic B cells. (**A**) B-cell frequency as a percentage of total immune cells in active and torpor groups. B-cell frequency was significantly reduced in the torpor group compared with active controls. Data are presented as mean ± SEM, with *n* = 4 animals per group; each dot represents one animal. Statistical significance was determined using an unpaired two-tailed Welch’s *t*-test. \**P* < 0.05. (**B**) UMAP visualization of B cells within the integrated dataset (right), with cells from active (left, red) and torpor (middle, blue) animals projected onto the same embedding. (**C**) The volcano plot shows gene expression differences between torpid and active animals. Red points indicate genes with nominal *P* < 0.05 and |log₂ fold change| > 2. Selected genes are labeled. (**D**) Heatmap of the top 30 differentially expressed genes in B cells between active and torpor groups. Colors indicate scaled gene expression across samples. (**E**) IPA canonical pathways in torpid versus active animals. Twenty representative pathways meeting both criteria (Benjamini–Hochberg-adjusted *P* < 0.05 and |z-score| ≥ 2) are shown. Bar length represents the absolute IPA activation z-score. Red and blue indicate predicted activation and inhibition, respectively. The dashed line marks |z-score| = 2.

Differential expression analysis identified extensive transcriptional remodeling in B cells during torpor compared with those from active animals. Of the 18,452 genes detected, 1,340 were differentially expressed genes (DEGs) (adjusted *P* < 0.05 and |log₂ fold change| > 1), of which 610 were upregulated and 730 downregulated during torpor (Supplementary Table 1). This 52.4% reduction in B cell frequency was accompanied by a coherent transcriptional shift away from activation. Of the top down-regulated DEGs, *Skap2* plays a critical role in BCR-mediated signaling, and *Skap2* knockout B cells exhibit significantly reduced BCR-mediated proliferation and activation^44,45^; *Tlr3* is expressed in a subset of B cells and CD138+ plasma cells, recognizing double-stranded RNA, activating IRF3 and NF-κB pathways, and promotes B cell cytokine secretion, costimulatory molecule up-regulation, isotype class switching, and antibody production; *Ints4* encodes a core component of the Integrator complex that regulates RNA polymerase II transcription termination and RNA processing, which is positively associated with B cell function^46^. Of the up-regulated DEGs, notably, *Txnip*, a key inhibitor of glucose uptake and glycolysis, was significantly upregulated (∼6.38-fold) and showed consistent expression across all torpid replicates (Fig. 3C-3D, Supplementary Table 1). Together with lower *Hk2* expression, higher *Txnip* expression suggests reduced glycolysis in splenic B cells during torpor. In addition, *Cirbp*, which encodes a cold-inducible RNA-binding protein, was more than 2-fold higher in torpid animals in all five cell types examined (2.1- to 3.5-fold; adjusted *P* < 0.05 in B cells, T cells, macrophages, and DCs; Supplementary Table 1), consistent with a cold-associated transcriptional response during torpor^47^. In B cells from torpid animals, IPA suggested inhibition of 12 of the 20 pathways shown in Fig. 3E. These included signaling pathways involved in B-cell activation and growth (ERK/MAPK, mTOR, STAT3, IL-2, and IGF-1 signaling) and in leukocyte trafficking (chemokine signaling and leukocyte extravasation signaling), as well as mitochondrial translation, asparagine N-linked glycosylation, regulation of eIF4 and p70S6K signaling, and trafficking and processing of endosomal Toll-like receptors. PFKFB4 signaling was also among the inhibited pathways, consistent with the lower *Hk2* and higher *Txnip* expression observed in B cells from torpid animals. IPA suggested activation of the remaining eight pathways, which included processing of capped intron-containing pre-mRNA, RNA polymerase II transcription, selected cell-cycle checkpoint pathways, and MHC class I–mediated antigen processing and presentation. Overall, torpor was associated with a lower relative frequency of splenic B cells, and the pathway analysis suggested suppression of multiple signaling pathways involved in B-cell activation, growth, and trafficking.

### Transcriptomic Reprogramming of Splenic T Cells during Torpor

Consistent with the significant reductions in CD3+ pan–T cells and CD4+ T cells detected using a combination of IHCS and QIA, scRNA-seq showed a lower relative frequency of splenic T cells in torpid animals than in active animals; however, this difference did not reach statistical significance (*P* = 0.480; Fig. 4A). Nevertheless, UMAP visualization revealed a pronounced transcriptional divergence, with torpid T cells forming a distinct cluster separate from active counterparts (Fig. 4B). Differential expression analysis identified a coherent torpor-associated T-cell program characterized by induction of metabolic inhibitors and repression of proliferative signaling (Fig. 4C). T cells displayed widespread transcriptional changes (1,163 DEGs), underscoring comprehensive reprogramming. This program was marked by the induction of key inhibitors, including the cell-cycle arrest gene *Cdkn1a* (fold change [FC] = 9.29) and the metabolic regulator *Txnip* (FC = 4.86), whereas *Hk2* expression was lower (Supplementary Table 1). The splenic T cells from torpid animals were characterized by higher expression of *Cdkn1a* (encoding p21), *Cdkn1b* (encoding p27), the antiproliferative immediate-early gene *Btg2*, the metabolic regulator *Txnip*, and the Notch target gene *Hes1* (FC = 12.67) (Fig. 4C). Expression of proliferation-associated genes (e.g., *Tk1*, *Kif18a*) was concurrently suppressed. Hierarchical clustering of the top 30 DEGs confirmed that this transcriptional shift was highly consistent across biological replicates (Fig. 4D).

**Figure 4.**
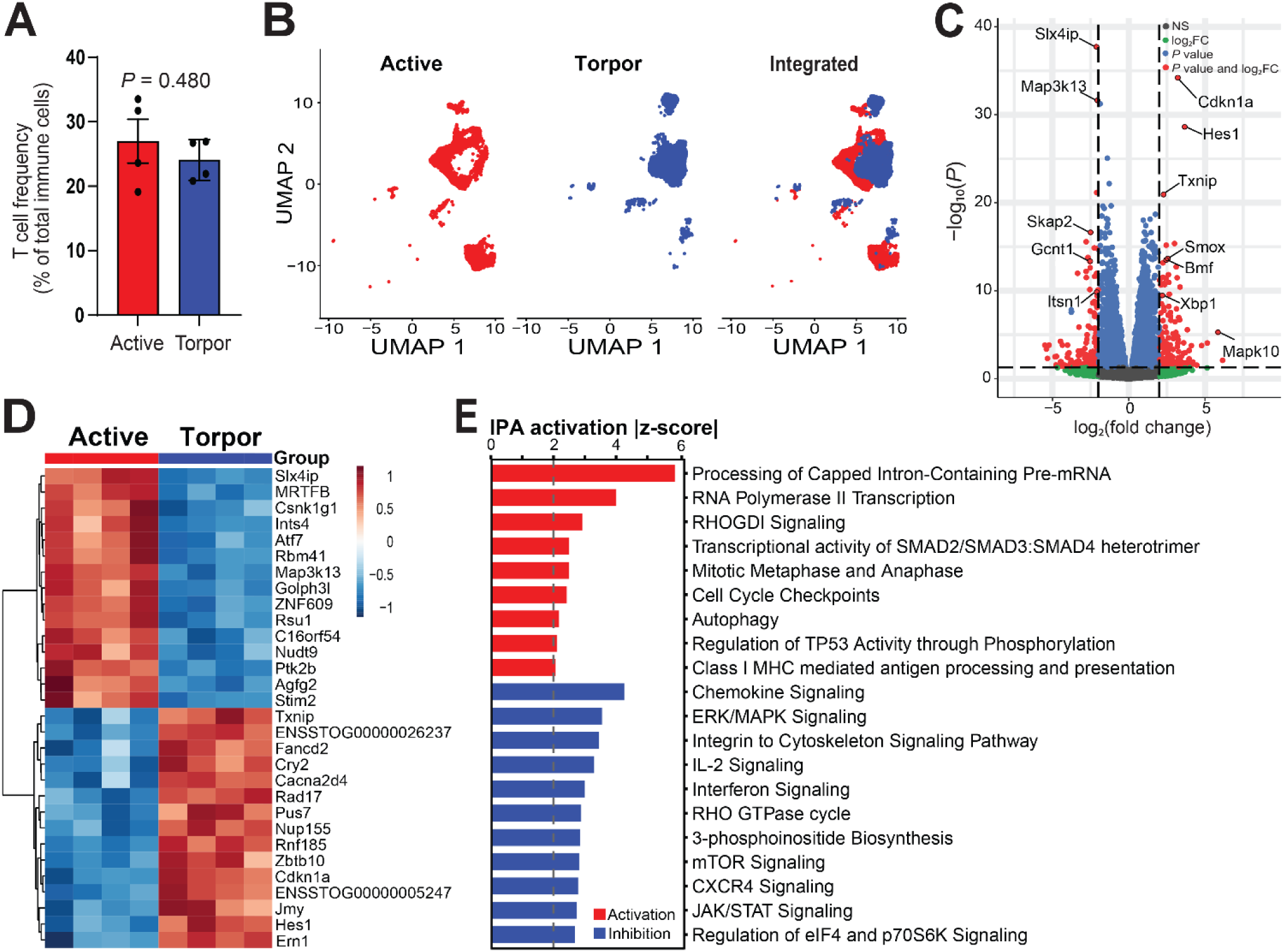
Torpor alters the transcriptional profile of splenic T cells. (**A**) T-cell frequency as a percentage of total immune cells in active and torpor groups. No significant difference in overall T-cell frequency was observed between the groups. Data are presented as mean ± SEM, with *n* = 4 animals per group; each dot represents one animal. Statistical significance was assessed using an unpaired two-tailed Welch’s *t*-test. (**B**) UMAP visualization of T cells within the integrated dataset (right), with cells from active (left, red) and torpor (middle, blue) animals projected onto the same embedding. (**C**) The volcano plot shows gene expression differences between torpid and active animals. Red points indicate genes with nominal *P* < 0.05 and |log₂ fold change| > 2, with selected representative genes labeled. (**D**) Heatmap of the top 30 differentially expressed genes in T cells between active and torpor groups. Colors indicate scaled gene expression across samples. (**E**) IPA canonical pathways in torpid versus active animals. Twenty representative pathways meeting both criteria (Benjamini–Hochberg-adjusted *P* < 0.05 and |z-score| ≥ 2) are shown. Bar length represents the absolute IPA activation z-score. Red and blue indicate predicted activation and inhibition, respectively. The dashed line marks |z-score| = 2.

In splenic T cells from torpid animals, IPA predicted inhibition of 11 of the 20 pathways shown in Fig. 4E. These included pathways involved in T-cell activation and cytokine responses (ERK/MAPK, IL-2, JAK/STAT, interferon, and mTOR signaling) and in cell migration and cytoskeletal regulation (chemokine signaling, CXCR4 signaling, integrin-to-cytoskeleton signaling, and the RHO GTPase cycle), as well as regulation of eIF4 and p70S6K signaling. Together with the higher expression of cell-cycle inhibitors described above, these findings suggest suppression of multiple pathways involved in T-cell activation, cytokine responses, and migration during torpor. IPA predicted activation of the remaining nine pathways, including capped intron-containing pre-mRNA processing, RNA polymerase II transcription, autophagy, TP53 regulation by phosphorylation, and selected cell-cycle pathways. Together with the gene expression changes described above, these findings are consistent with a more quiescent transcriptional profile in splenic T cells during torpor.

### Increased Relative Frequency and Altered Transcriptional Profiles of Splenic Macrophages

Macrophages exhibited a significant increase in relative abundance during torpor (*P* = 0.017; Fig. 5A). Integrated UMAP analysis demonstrated clear segregation of macrophages by torpid and active groups, with torpid macrophages forming a distinct transcriptional cluster (Fig. 5B). This enrichment suggests preferential retention of macrophages within the spleen during torpor. Macrophage gene expression analysis identified 222 DEGs from torpid versus active animals (adjusted *P* < 0.05 and |log₂FC| > 1), including 69 upregulated and 153 downregulated during torpor (Supplementary Table 1). Torpid macrophages upregulated lipid metabolism genes (*Ldlr*, *Sqle*), suggesting a functional retooling to handle altered lipid profiles during prolonged fasting, pivoting from inflammation to homeostasis. Hierarchical clustering confirmed consistent expression patterns across all biological replicates (Fig. 5D). Canonical pathway analysis using IPA identified the inhibition of pathways involving cytoskeletal regulation, including the RHO GTPase cycle, from torpid animals (Fig. 5E), which is related to cellular signaling and motility. Together with the increased relative frequency of macrophages, DEGs and altered canonical pathways demonstrate differences in cellular compositional and transcriptional profiles between torpid and active groups.

**Figure 5.**
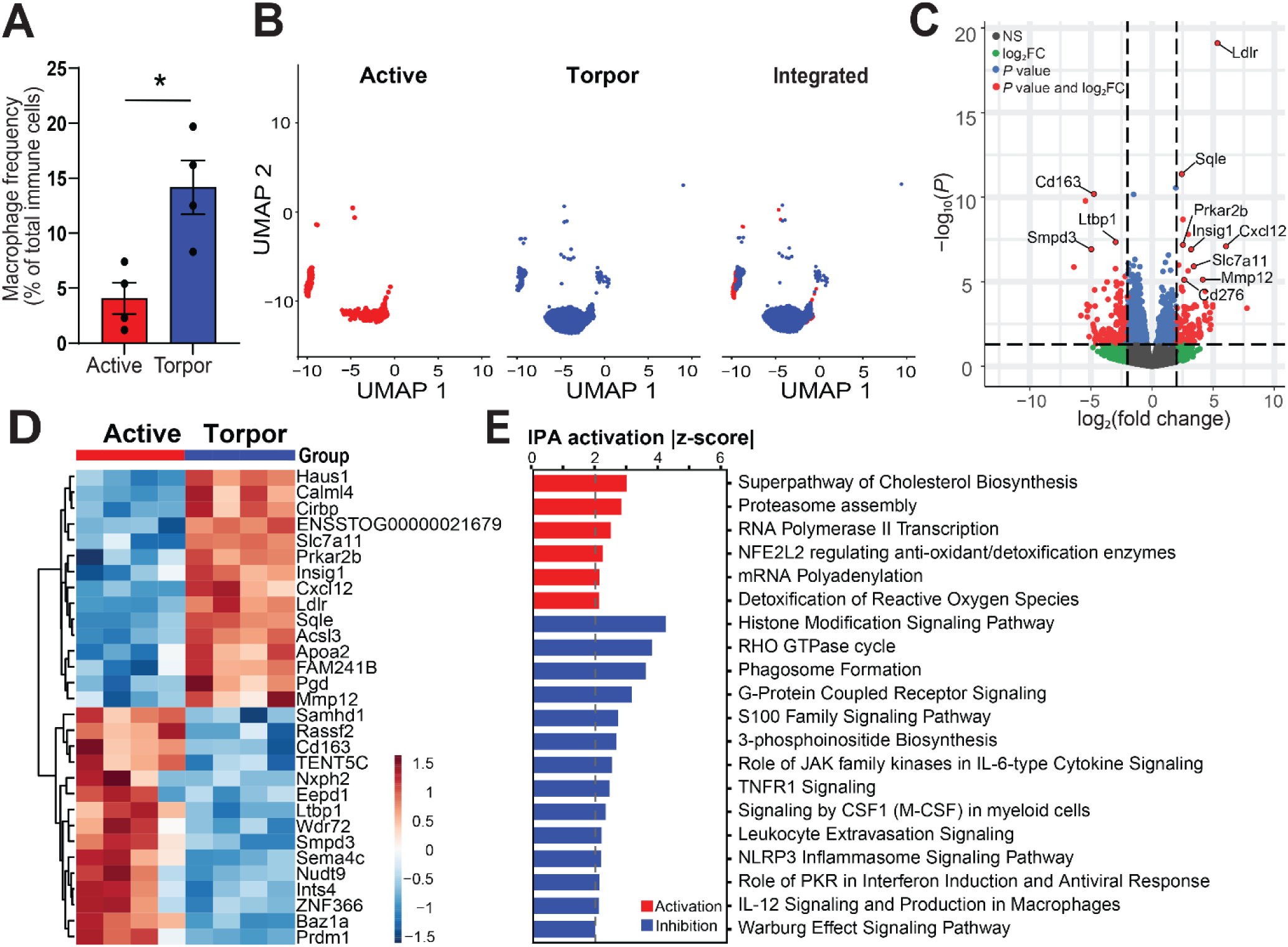
Torpor increases splenic macrophage relative abundance and alters their transcriptional profile. (**A**) Macrophage frequency as a percentage of total immune cells in active and torpor groups. Macrophage frequency was significantly increased in the torpor group compared with active controls. Data are presented as mean ± SEM; with *n* = 4 animals per group, each dot represents one animal. Statistical significance was determined using an unpaired two-tailed Welch’s *t*-test. \**P* < 0.05. (**B**) UMAP visualization of macrophages within the integrated dataset (right), with cells from active (left, red) and torpor (middle, blue) animals projected onto the same embedding. (**C**) Volcano plot of gene expression in macrophages from torpor versus active animals. Genes meeting nominal *P* < 0.05 and |log₂ fold change| > 2 are highlighted in red, with selected representative genes labeled. (**D**) Heatmap of the top 30 differentially expressed genes in macrophages between active and torpor groups. Colors indicate scaled gene expression across samples. (**E**) IPA canonical pathways in torpid relative to active animals. Shown are 20 representative pathways with Benjamini–Hochberg-adjusted *P* < 0.05 and |z-score| ≥ 2. Bar length, absolute activation z-score; red, predicted activation; blue, predicted inhibition; dashed line, |z-score| = 2.

### Torpor-Associated Suppression of Innate Immune Signaling in Splenic DCs

DCs showed a higher relative frequency in torpid animals, although the difference did not reach statistical significance (*P* = 0.059, Fig. 6A). Their transcriptional profiles differed between the active and torpid groups (Fig. 6B). Five genes met the joint differential-expression criteria of adjusted *P* < 0.05 and |log₂ fold change| > 1, and all five are displayed in the heatmap (Fig. 6C, D). Of these, *Cirbp* showed higher expression in torpid animals, whereas *Cdcp1*, *Arhgap22*, *Nudt9*, and *Hipk2* showed lower expression. Using the separate input criteria described in the Methods, IPA predicted inhibition of two innate immune pathways in DCs from torpid animals: inflammasome signaling and DDX58/IFIH1-mediated induction of interferon-α/β (Fig. 6E). IPA also predicted inhibition of PDGF and HGF signaling, alongside activation of pathways involved in DNA synthesis and mitotic metaphase and anaphase (Fig. 6E).

**Figure 6.**
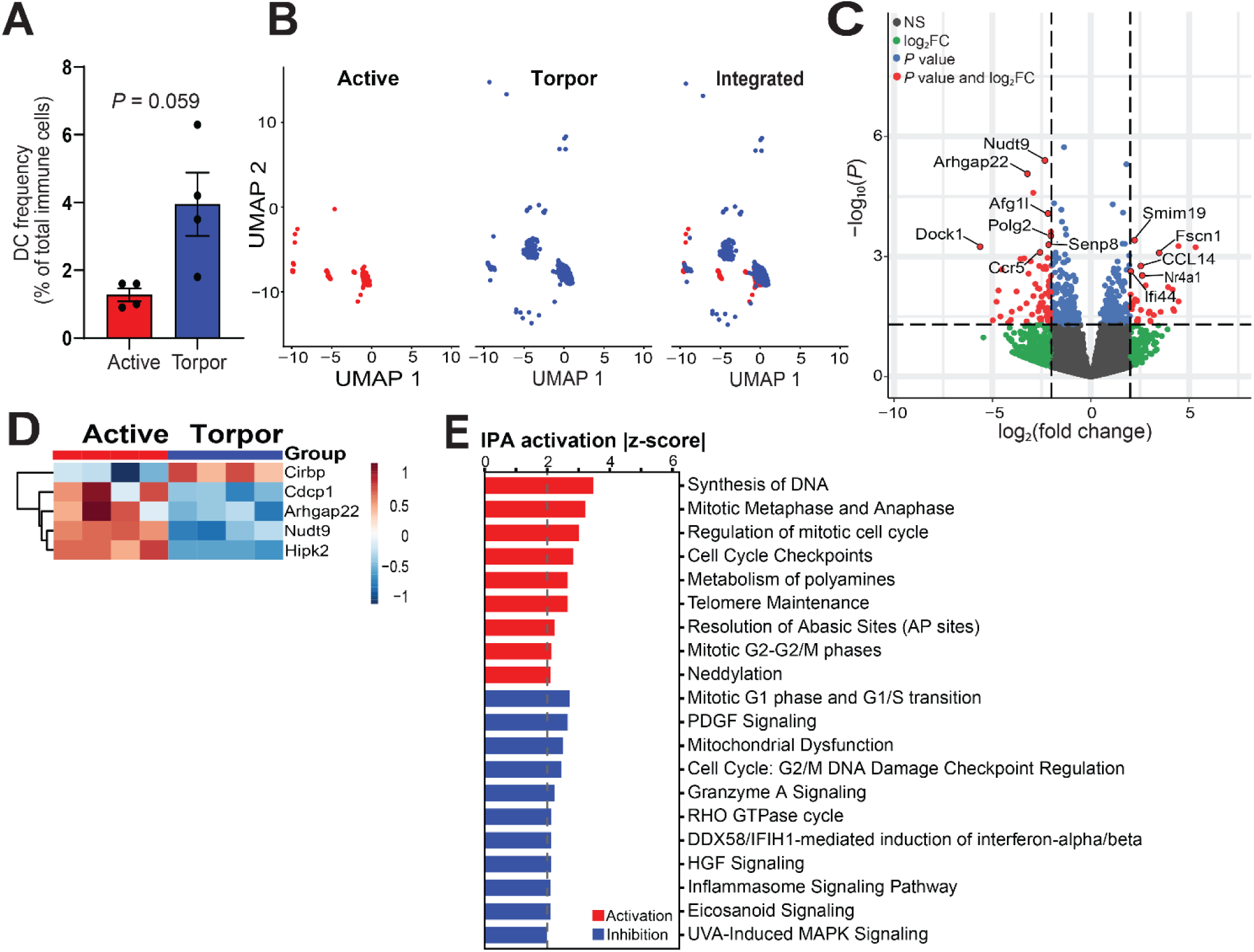
Torpor-associated transcriptional changes in splenic dendritic cells. (**A**) Relative frequency of dendritic cells (DCs), expressed as a percentage of total immune cells, in active and torpid 13-LGS. Each point represents one animal, with *n* = 4 animals per group, and bars show mean ± SEM. Statistical significance was assessed using an unpaired two-sided Welch’s *t*-test (*P* = 0.059). (**B**) Uniform manifold approximation and projection (UMAP) visualization of DCs from active animals (red), torpid animals (blue), and the integrated dataset. (**C**) Volcano plot of gene expression in DCs from torpid relative to active animals. Positive and negative log₂(fold change) values indicate genes upregulated and downregulated during torpor, respectively. Genes meeting only the fold-change threshold are shown in green, those meeting only the *P*-value threshold in blue, and those meeting both thresholds in red; nonsignificant genes are shown in gray. Dashed lines indicate the predefined fold-change and statistical-significance thresholds. Selected genes are labeled. (**D**) Heatmap showing scaled expression of selected DEGs in DCs from active and torpid animals. Columns represent individual animals, and rows represent genes. Colors indicate row-scaled relative expression, with red and blue denoting higher and lower expression, respectively. (**E**) IPA canonical pathways in torpid versus active animals. Twenty representative pathways meeting both criteria (Benjamini–Hochberg-adjusted *P* < 0.05 and |z-score| ≥ 2) are shown. Bar length represents the absolute IPA activation z-score. Red and blue indicate predicted activation and inhibition, respectively. The dashed line marks |z-score| = 2.

### Torpor-Associated Transcriptional Changes in Splenic Neutrophils

Neutrophils showed a trend toward increased abundance during torpor (Fig. 7A) and formed transcriptionally distinct clusters between torpor and active groups (Fig. 7B). In neutrophils, 78 genes met both criteria (adjusted *P* < 0.05 and |log₂FC| > 1), including 43 upregulated and 35 downregulated genes in torpid versus active animals (Supplementary Table 1); the top DEGs are displayed in the volcano plot and heatmap (Fig. 7C, D). Compared with those from active animals, splenic neutrophils from torpid animals showed lower expression of *Syk*, *Hk2* (encoding the glycolytic enzyme hexokinase 2), and the mitosis-associated genes *Kif11* and *Ccnb2* (Fig. 7C, D). These cells also showed higher expression of *Il1rap*, which encodes the IL-1 receptor accessory protein, as well as *Nfkbia* and *CXCL8* (Fig. 7C, D). Consistent with these expression patterns, IPA predicted inhibition of HIF-1α signaling and JAK1/JAK3-dependent γc cytokine signaling, together with activation of IL-1 family signaling (Fig. 7E).

**Figure 7.**
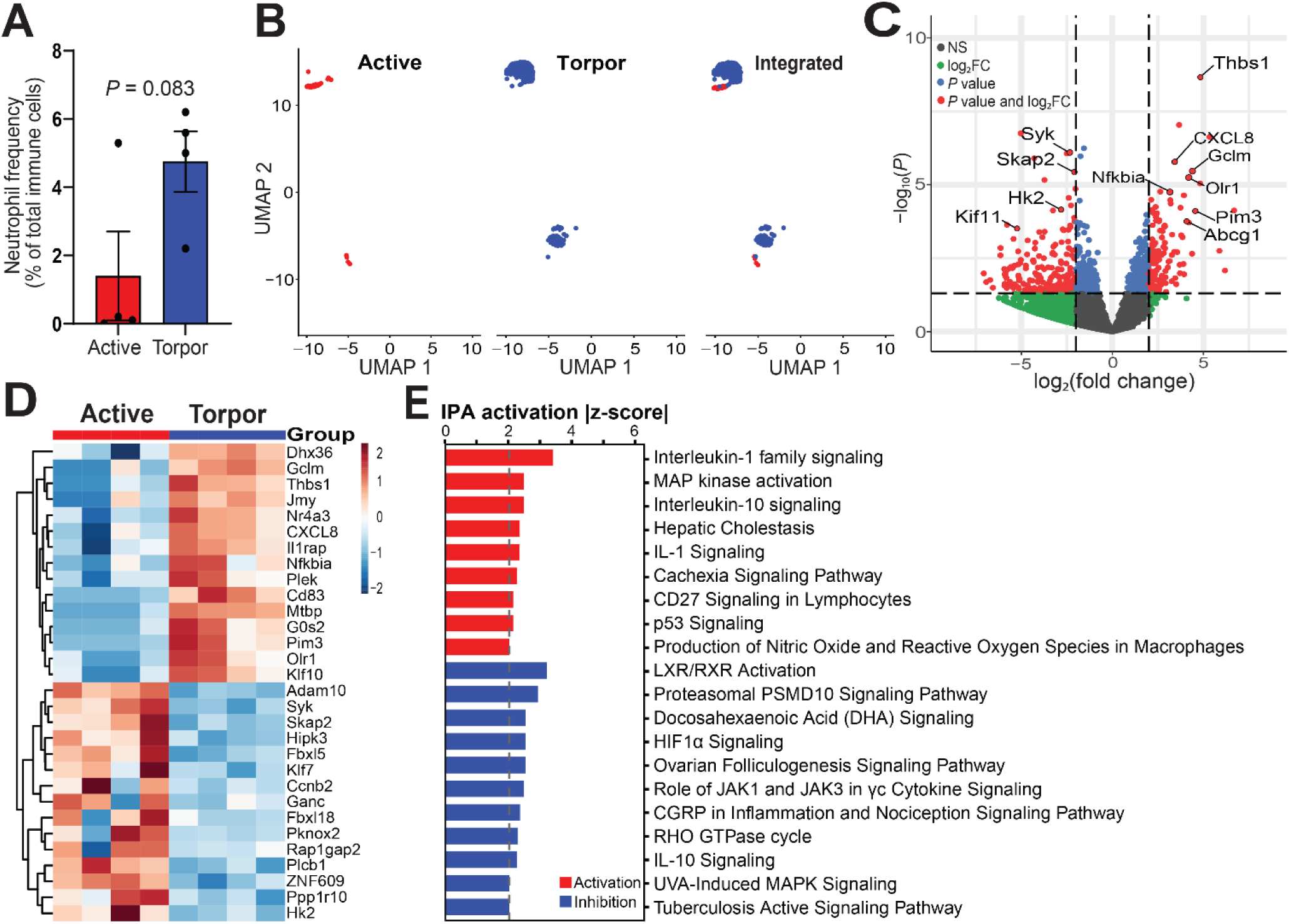
Torpor alters the transcriptional profile of splenic neutrophils. (A) Neutrophil frequency as a percentage of total immune cells in active and torpor groups. Data are presented as mean ± SEM; each dot represents one animal (*n* = 4 per group). No significant difference was observed between groups. Statistical significance was assessed using Welch’s unpaired *t*-test (*P* = 0.083). (B) UMAP visualization of neutrophils within the integrated dataset (right), with cells from active (left, red), and torpor (middle, blue) animals projected onto the same embedding. (**C**) The volcano plot shows gene expression differences in neutrophils between torpid and active animals. Red points indicate genes with nominal *P* < 0.05 and |log₂ fold change| > 2. Selected genes are labeled. (**D**) Heatmap of the top 30 differentially expressed genes in neutrophils between active and torpor groups. Colors indicate scaled gene expression across samples. (**E**) IPA canonical pathways in torpid versus active animals. Twenty representative pathways meeting both criteria (Benjamini–Hochberg-adjusted *P* < 0.05 and |z-score| ≥ 2) are shown. Bar length represents the absolute IPA activation z-score. Red and blue indicate predicted activation and inhibition, respectively. The dashed line marks |z-score| = 2.

## Discussion

Thirteen-lined ground squirrels can safely transition between torpor and arousal during hibernation and resume seasonal activity in spring^6,48^. During torpor, 13-LGS undergo marked metabolic depression accompanied by substantial suppression of immune-cell abundance and function. These changes reverse during the active season, when immune-cell populations and functional responsiveness are restored^20,22,23^. This physiological cycle provides a unique opportunity to investigate how secondary lymphoid tissues—particularly the spleen—adapt to extreme metabolic conditions at the tissue, cellular, and molecular levels. Previous work has established that torpor is accompanied by profound immune suppression, including marked blood leukopenia^22,23^, reduced serum complement activity^24^, no febrile response to bacterial LPS^20^, reduced humoral immune response to immunizations^25^, and suppressed cellular immune response in skin allografts during hibernation^26^. However, previous studies in hibernating mammals have not resolved immune cell lineage-specific states or spatial remodeling within LTs. Moreover, previous transcriptomic analyses of hibernating mammals conducted using microarrays and bulk mRNA-seq, have revealed gene expression programs associated with torpor in skeletal muscle, heart, liver, adipose tissue, bone marrow, and peripheral blood mononuclear cells. Yet the LTs, including the spleen, have not been fully studied. To fill this gap, in this study, we intended to better understand how immune suppression during torpor is implemented in the spleen at organ, tissue, cell, and molecular levels. We performed a comparative analysis of splenic architecture and immune cell composition between torpid and active 13-LGS using scRNA-seq combined with immunohistochemical staining (IHCS) and quantitative image analysis (QIA) of immune cells in the spleen.

We found that despite the profound metabolic depression in torpor, total spleen mass and spleen index were not significantly different between torpid and active 13-LGS (Fig. 1A), providing no evidence of global splenic atrophy. Instead, tissue staining/IHCS and QIA revealed a selective remodeling of splenic architecture characterized by a significantly reduced white-pulp area fraction (12.53 ± 1.05% in torpor versus 20.26 ± 2.10% in active control) with a relative expansion of red-pulp (Fig. 1B; *P* < 0.05). The white-pulp encompasses T cell–dense periarteriolar lymphoid sheaths and B cell-rich follicles. The reductions in PNA+ B cells and CD21+ cells (including B cells and follicular dendritic cells), and CD3+ and CD4+ T cells provide additional evidence of reduced white-pulp fraction and altered splenic tissue organization during torpor (Fig. 1C–E), likely reflecting a selective suppression of adaptive immune architecture in LTs and diminished lymphocyte proliferation and activation (Fig. 1F). These observations are consistent with the attenuated humoral and cellular adaptive immune responses during torpor^25,26^. The observed changes in splenic adaptive immune compartments may have implications for T-cell-dependent humoral responses, although their functional consequences remain to be established. The red-pulp functions as a blood filter and reservoir for blood immune cells^41,42^. Circulating leukocyte counts fall during torpor and recover upon arousal^22,23^. The relative expansion of red-pulp during torpor is consistent with the hypothesis that circulating leukocytes sequester within the spleen during torpor, thereby conserving energy while maintaining the capacity for rapid reconstitution of blood immune cells upon arousal. However, a relative increased ratio of red-pulp relative to white-pulp does not directly demonstrate the leukocyte sequestration in torpid spleen. Future study is needed to simultaneously quantify leukocyte numbers in the splenic red-pulp and peripheral blood compartments to assess their distribution in torpid and aroused animals.

Single-cell RNA sequencing (scRNA-seq) identified both compositional and transcriptional differences in splenic immune cells from torpid versus active 13-LGS. Consistent with the results of significantly lower frequency of B and T cells in torpid animals as compared with active animals using IHCS and QIA, scRNA-seq also identified a significant reduction of B cell frequency, where the T-cell frequency was reduced in torpid animals, but was not statistically significant. Despite these compositional differences, B and T cells shared several altered canonical pathways, including inhibition of ERK/MAPK, mTOR, and chemokine signaling and activation of capped intron-containing pre-mRNA processing and RNA polymerase II transcription (Figs. 3E and 4E). Thus, the lymphocyte response was not characterized by uniform suppression of transcription-associated processes. These results support selective remodeling of lymphocyte transcriptional programs. Nevertheless, they do not establish preservation of absolute T-cell numbers or enhanced survival, and the absence of activated apoptosis-related pathways cannot exclude cell loss.

Splenic myeloid populations also showed distinct responses. Macrophages had a significantly higher relative frequency in torpid animals, whereas the higher DC frequency did not reach statistical significance. Macrophage pathway predictions included inhibition of cytoskeletal regulatory programs, while DCs showed activation of selected cell-cycle-related pathways together with inhibition of inflammasome signaling. Neutrophils exhibited predicted activation of IL-1-related pathways, whereas two IL-10-related pathway entries showed opposite predicted directions (Figs. 5E–7E). These differences argue against describing all innate immune populations as uniformly suppressed during torpor. Instead, they indicate cell-type-specific transcriptional changes whose consequences for migration, inflammatory responses, and host defense require functional assessment. In golden-mantled ground squirrels, Maniero found that splenic macrophages collected at different stages of the hibernation cycle could bind LPS in vitro even at low temperatures^49^.

Metabolic and proliferation-associated findings likewise require interpretation at the level measured. Reduced expression of selected proliferation-associated genes, together with the lower Ki67-positive staining-area fraction, was consistent with reduced proliferation-associated activity in torpid spleens. However, the IPA results did not indicate uniform inhibition of all cell-cycle, RNA-processing, or biosynthetic pathways. Changes in metabolic-regulator transcripts identify candidate processes for further investigation but do not directly measure glucose uptake, substrate availability, or glycolytic flux. Similarly, a lack of consistent transcriptional induction of fatty-acid oxidation genes does not exclude altered lipid utilization. The present data therefore support changes in metabolic regulation at the transcript level without establishing a universal reduction in cellular energy throughput or a specific fuel-switching mechanism.

A notable finding was the predicted activation of capped intron-containing pre-mRNA processing in both B and T cells. This result contrasts with a model of uniformly suppressed RNA-related programs during torpor and instead indicates selective changes in the expression of RNA-processing machinery. Increased abundance of these transcripts does not necessarily imply increased RNA-processing rates at the low body temperatures characteristic of torpor. It also does not establish increased global transcription or protein synthesis. One possibility is that altered RNA-processing programs contribute to transcript maintenance or remodeling during torpor; however, this remains a hypothesis. Measurements of RNA processing, transcript stability, and protein synthesis across torpor and arousal would be needed to determine the functional significance of this signature. Together, the tissue and single-cell findings support a model of selective splenic remodeling associated with torpor. Changes in tissue organization and immune-cell representation were accompanied by distinct transcriptional responses across lymphoid and myeloid populations. Shared pathway predictions in B and T cells coexisted with differences among macrophages, DCs, and neutrophils. These observations do not establish a hierarchical regulatory mechanism or demonstrate that the changes conserve energy, preserve immune surveillance, or prepare cells for rapid recovery. Temperature-associated, endocrine, and seasonal influences are possible contributors, but their individual roles cannot be resolved by the present comparison.

This study has several limitations. First, active animals were sampled in September, whereas torpid animals were sampled in December–January under hibernation-inducing conditions; therefore seasonal and housing effects cannot be excluded. Second, immune cell abundance measured by IHCS and QIA or scRNA-seq reflects relative cell frequencies rather than absolute cell numbers. Third, spleen weight, length, and splenic index did not differ significantly between groups, but these measurements do not exclude splenic atrophy. Finally, the absence of interbout-arousal or recovery groups and functional perturbation experiments limits conclusions about reversibility and causality. Future studies combining absolute cell quantification, functional assays, and sampling across the hibernation cycle will be needed to resolve these questions.

In conclusion, torpid versus active 13-LGS differed in splenic architecture, relative immune-cell composition, and cell-type-specific transcriptional profiles. These findings identify selective immune remodeling associated with torpor and provide a basis for investigating its functional consequences across the hibernation cycle.

## Acknowledgements

We thank Dirk Anderson, Director of the Biotechnology’s Flow Cytometry & Single-Cell Genomics Core Facility at the University of Nebraska–Lincoln (Morrison Center, Room 160), for assistance with single-cell RNA-sequencing library preparation. We also thank the staff of the Life Sciences Annex (LSCA) at the University of Nebraska–Lincoln for their support with the animal experiments. We are grateful to K. M. Main Uddin and Dr. Yilun Cheng of Dr. Q. Li’s laboratory, as well as Frazer I. Heinis of Dr. Matthew T. Andrews’s laboratory, for their assistance with sample preparation and collection. M.T.A. acknowledges support from the University of Nebraska Institute of Agriculture and Natural Resources.

## Author contributions

Drs. QL, MZ, and MA conceived the idea. MZ and QL wrote the manuscript. MZ conducted all experiments and data analysis. MZ and RB collected the samples, while MZ, JC, RB, SM, and SCL contributed to sample processing and project discussions. Drs. JW, WY, and CZ contributed to the scRNA-seq analysis. All authors discussed the results and revised the manuscript.

## Data availability

The single-cell RNA-sequencing data generated in this study have been deposited in the NCBI Gene Expression Omnibus (GEO) under accession code GSE347994. All other data supporting the findings of this study are available within the Article and its Supplementary Information.

