## Supplementary Figures for "Hibernation-Driven Immune Suppression in Thirteen-Lined Ground Squirrels": Supplementary Materials--JI.pdf

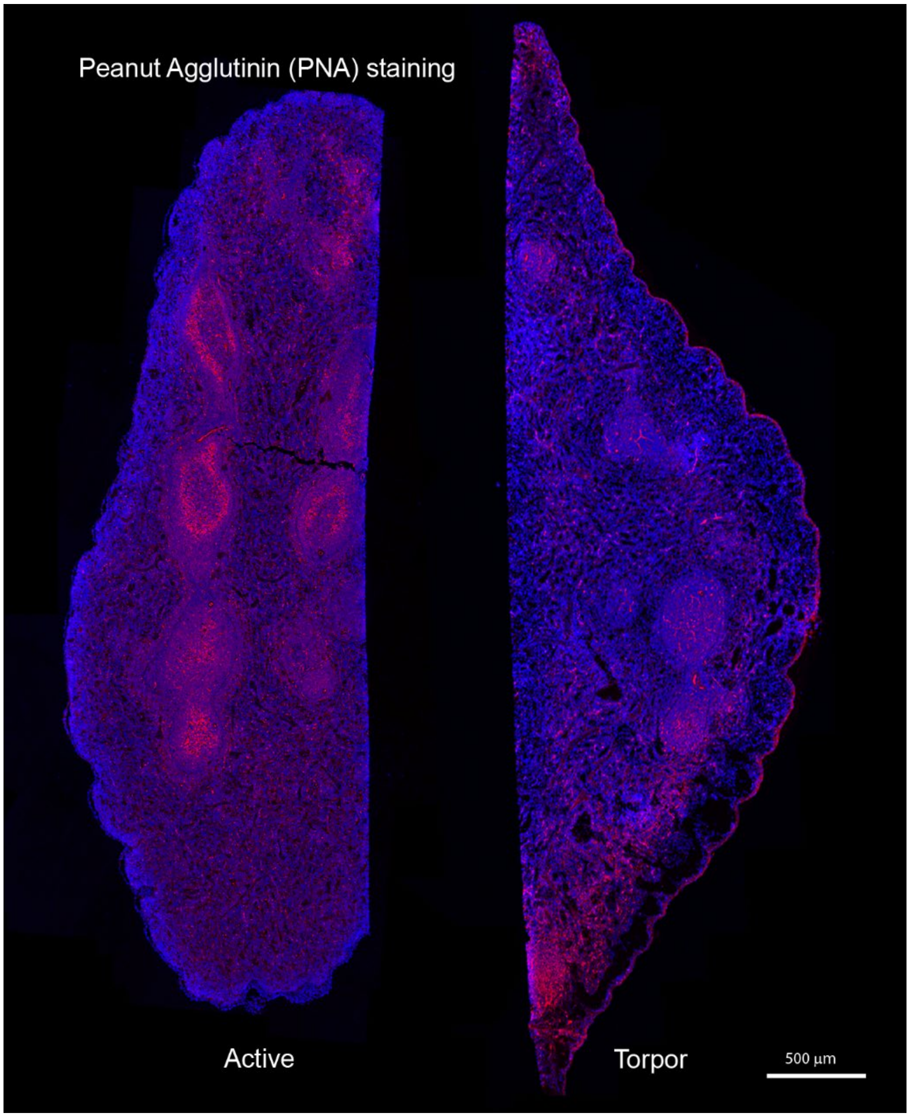


**Supplementary Figure 1. Peanut agglutinin (PNA) staining of splenic germinal-center B cells in active and torpid 13-lined ground squirrels.**

Representative whole-section fluorescence images of spleens from active and torpid animals following neuraminidase treatment and staining with Alexa Fluor 647-conjugated PNA, PNA fluorescence is shown in red, and nuclei counterstained with DAPI are shown in blue. PNA-positive germinal-center B cells were readily detectable in active spleens but markedly diminished during torpor. Images are representative of the spleen sections examined in each physiological group. Scale bars, 500 µm.

**Supplementary Table 1. Cell type–resolved differential gene expression in splenic immune cells between active and torpid 13-lined ground squirrels**
